# A sequential event-responsive fluorescent reporter for inducible Cre and Flp combinatorial recombination

**DOI:** 10.1101/2024.11.20.624546

**Authors:** Giada Pessina, Mattia Camera, Fabrizio Loiacono, Antonio Uccelli, Fabio Benfenati, Paolo Medini, Francesco Trovato, Sebastian Sulis Sato

## Abstract

Current technologies for precise genetic manipulation of cells often utilize site-specific recombinases, enabling the creation of conditional transgenic models for studying gene functions in specific tissues and at defined time windows. The advent of drug-inducible recombinases has further enhanced this field by allowing temporal control of gene expression. However, detecting gene expression patterns when multiple recombinases are used is challenging, particularly when visualizing combinatorial maps and their temporal profile. To address this, we developed Rubik, a fluorescent genetic reporter specifically designed to monitor the action of Cre and Flp recombinases by expressing different fluorophores depending on the temporal sequence of recombination events. Rubik offers four alternative fluorescent configurations: blue for Cre recombination, green for Flp recombination, yellow for Cre followed by Flp recombination, and red for Flp followed by Cre recombination. Additionally, Rubik integrates light-gated ion channels, allowing optogenetic excitation or inhibition of electrically-excitable cells based on the temporal sequence of recombination events. Moreover, to add temporal control over Rubik recombination, we optimized a trimethoprim-inducible FlpO recombinase (FlpO-DD) to use in combination with the tamoxifen-inducible Cre recombinase (ERT2CreERT2). We validated Rubik by generating a stable knock-in HeLa cell line using CRISPR-Cas9 and single-cell sorting. Our results confirmed the effective functioning of Rubik with Cre and Flp recombinases and showed the ability to induce selective recombination via either tamoxifen or trimethoprim. This system holds potential for neuroscience research, particularly in generating knock-in mice expressing Rubik, which could be instrumental in precisely defining the roles of specific neuronal populations based on Cre and Flp activity.

## Introduction

Site-specific recombination (SSR) is one of the most widely used molecular tools to control gene expression. Indeed, Cre-Lox conditional systems have been used to confine the expression of transgenes in specific cell types and tissues in a variety of animal models^1,2^. Further development of drug-inducible Cre recombinases^3–5^ allowed to exert temporal control over gene expression, giving rise to conditional-inducible models. This technology is necessary for genes that cannot be constitutively knocked out due to their essential role in development^6^. Moreover, conditionally inducible systems have been used together with activity-dependent gene expression systems to label and manipulate specific neuronal populations that are active during a memory recall or behavioral task^7–9^. Combinatorial recombination, as used in the Brainbow technique^10^, has been used to label neurons with multiple fluorescent reporters following randomized recombination sequences. Brainbow offers a powerful tool for gaining resolution in the study of neuronal anatomy. However, the randomization of recombination events can be a limitation, as the different combinatorial results are not associated with a functional or anatomical property of cells. More recently, the Cre-Lox system has been combined with the Flp-FRT system to create a molecular tool in which the outcome of recombination events is determined by Boolean logic operations^11^. Among the Boolean logic operators implemented, the “AND” operator (Cre AND Flp recombination) is especially relevant when studying brain connectivity. Indeed, this tool has been used in combination with anterograde transsynaptic viral vectors to trace and manipulate specific anatomical convergence in the mouse brain: a neuron expresses a fluorescent reporter and an optogenetic tool only if it receives a direct connection from two different brain regions^12^. However, this approach is still limited by the fact that only neurons expressing both Cre and Flp recombinases can be tagged. Different recombination strategies have been developed to differentially label also recombination by Cre and not Flp, or vice versa (AND NOT Boolean operation)^11^. Nevertheless, none of these tools offers the possibility to perform both these Boolean operations simultaneously. Furthermore, when studying the function of specific neuronal populations, both optogenetic excitation and inhibition are often needed. In this work, we describe Rubik, a genetically encoded tool consisting of four different SSR-dependent fluorescent reporters that offers the possibility of simultaneously tagging multiple neuronal populations based on the sequential expression of Cre and Flp recombinases. Moreover, Rubik allows to perform both optogenetic excitation and inhibition experiments. These two conditions can be achieved by switching the order of recombination events: Cre followed by Flp (Cre→Flp) or Flp followed by Cre (Flp→Cre) result in two different configurations of Rubik, leading to the expression of either an excitatory or an inhibitory opsin^13,14^ together with a different fluorescent protein. Additionally, to have temporal control over recombination events, we combined Rubik with drug-inducible recombinases. Although tamoxifen-inducible Cre recombinase (ERT2CreERT2) has been widely used and tested^11^, an effective drug-inducible system for Flp with a non-tamoxifen-based drug is still missing. For this reason, we generated a new trimethoprim-inducible FlpO system (FlpO-DD) based on a Flp recombinase fused with an N-terminal destabilization domain^15^.

## Results

### Rubik’s design

Rubik is a fluorescent genetic reporter specifically designed to tag distinct cell populations based on the sequential action of Cre and Flp recombinases. Before recombination, no fluorescent reporter is expressed. Due to the presence of Cre-specific Lox sites and Flp-specific FRT sites, Rubik allows the expression of four distinct fluorescent proteins, depending on the sequence of recombination events: mTagBFP2 (blue) is expressed upon Cre, mTFP1 (green) upon Flp, mGold (yellow) upon sequential Cre→Flp, and mScarlet (red) upon sequential Flp→Cre recombination (**Figure 1**). Moreover, to allow optogenetic manipulation of tagged cells, the yellow and red fluorophores are associated with the co-expression of excitatory or inhibitory optogenetic tools, respectively. The optogenetic excitation is based on the light-gated ion channel ChR2, linked to the yellow fluorescent protein mGold through a P2A cleavage peptide, ensuring independent localization of the two proteins. The optogenetic inhibition is based on the light-gated anion channel GtACR1, linked through a T2A cleavage peptide to the red fluorescent protein mScarlet. Moreover, these opsins have a tag to confine their localization to the soma for more precise spatial localization of optogenetic stimulation. This setup allows to choose between photo-stimulation or photo-inhibition of tagged cells by controlling the temporal sequence of Cre and Flp activity.

**Figure 1.**
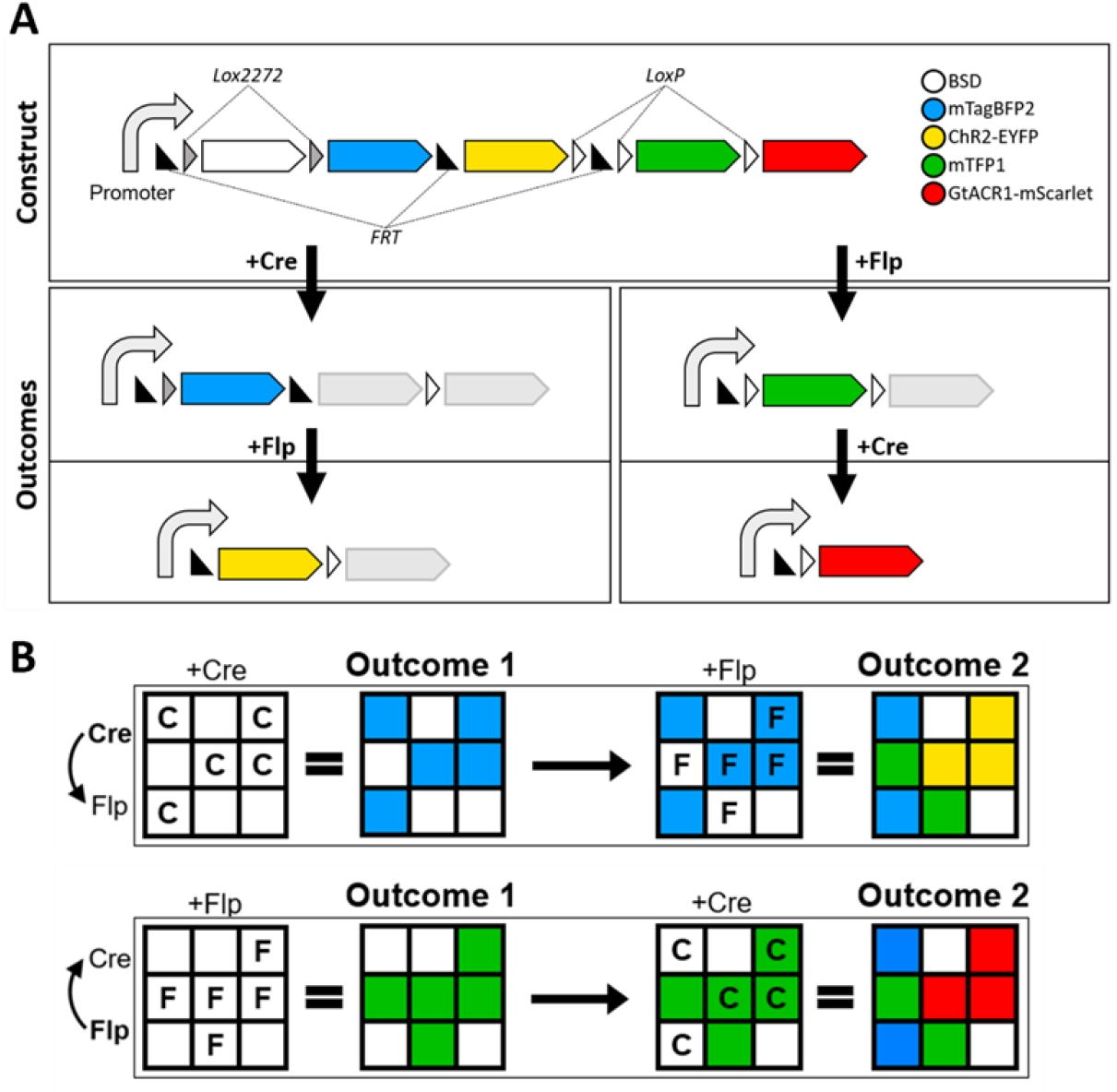
Design and functioning of Rubik. (A) Schematic representation of Rubik and its configurations following Cre and Flp recombination. Alongside fluorescent proteins, the Rubik construct includes a constitutive CAG promoter, BSD (blasticidin S deaminase), ChR2 (channelrhodopsin-2), and GtACR1-ST (Guillardia theta anion channelrhodopsin). Before recombination, no fluorescent reporter is expressed; sequential Cre/Flp recombination events activate different fluorescent reporters, as illustrated. (B) Rubik’s different outputs based on the order of recombination events: blue for single Cre recombination, green for single Flp recombination, yellow for sequential Cre→Flp recombination, and red for sequential Flp→Cre recombination.

### Validation of Rubik’s functioning

To validate and quantify the recombination efficiency and specificity of Cre and Flp recombination on the Rubik construct, we generated a monoclonal and stable HeLa cell line with Rubik integrated into the transcriptionally active site AAVS1 (Rubik-cells). We used CRISPR-Cas9, single-cell sorting, and antibiotic selection to obtain single clones of Rubik-cells, and we performed the validation following the schematic protocol reported in **Suppl. Figure 1**. Briefly, we transfected Rubik-cells with either Cre or Flp recombinases and sorted mTagBFP2^+^ (Rubik-Blue) and mTFP^+^ (Rubik-Green) cells (sorting strategies are shown in **Suppl. Figure 2C**), respectively. Then, we transfected Rubik-Blue cells with Flp and Rubik-Green cells with Cre and sorted mGold^+^ (Rubik-Yellow) and mScarlet^+^ (Rubik-Red) cells. We imaged sorted cells by confocal microscopy to obtain the emission spectra of each fluorescent reporter using four different excitation wavelengths (405, 458, 514, and 561 nm). Accordingly, we selected the combinations of excitation and emission wavelength ranges (see Material and Methods section) specific for each reporter (**Figure 2 A-D**). We evaluated the results of the four recombination by confocal microscopy using the spectral windows obtained before (**Figure 2 E-H**). We quantified the number of fluorescent cells for each condition by flow cytometry. Among Cre-transfected cells, about 33% were mTagBFP2^+^, while mTFP1^+^, mScarlet^+^, or mGold^+^ cell frequencies were below 0.2% (**Figure 3A**). Similarly, among Flp-transfected cells, about 45% were mTFP1^+^, about 2% were mGold^+^, while no mTagBFP2^+^ and mScarlet^+^ cell frequencies were detected (**Figure 3B**). Then, we sorted Rubik-Blue and Rubik-Green cells and transfected them with Flp and Cre recombinase, respectively. Among Flp-transfected Rubik-Blue cells, about 14% were mGold^+^, 52% were mTagBFP2^+^ (representing cells that did not undergo the second recombination), 5% were mTFP1^+^, and 30% were negative for all the fluorophores; mScarlet^+^ cell frequency was below 0.1% (**Figure 3C**). Similarly, among Cre-transfected Rubik-Green cells, about 9% were mScarlet^+^, 88% were mTFP1^+^ (representing cells that did not undergo the second recombination step), and 1% were negative for all the fluorophores; mTagBFP2^+^ and mGold^+^ cell frequencies were below 0.3%. (**Figure 3D**). The gating strategy and the negative control are shown in **Suppl. Figure 2 A, B**. Finally, we used sorted cells for each fluorophore to validate recombination events by PCR analysis, using primer pairs to specifically detect DNA sequences near the recombination sites involved in each recombination event (**Suppl. Figure 3**). These results show that Rubik is recombined by each recombinase as expected, with a negligible rate of non-specific recombination.

**Figure 2.**
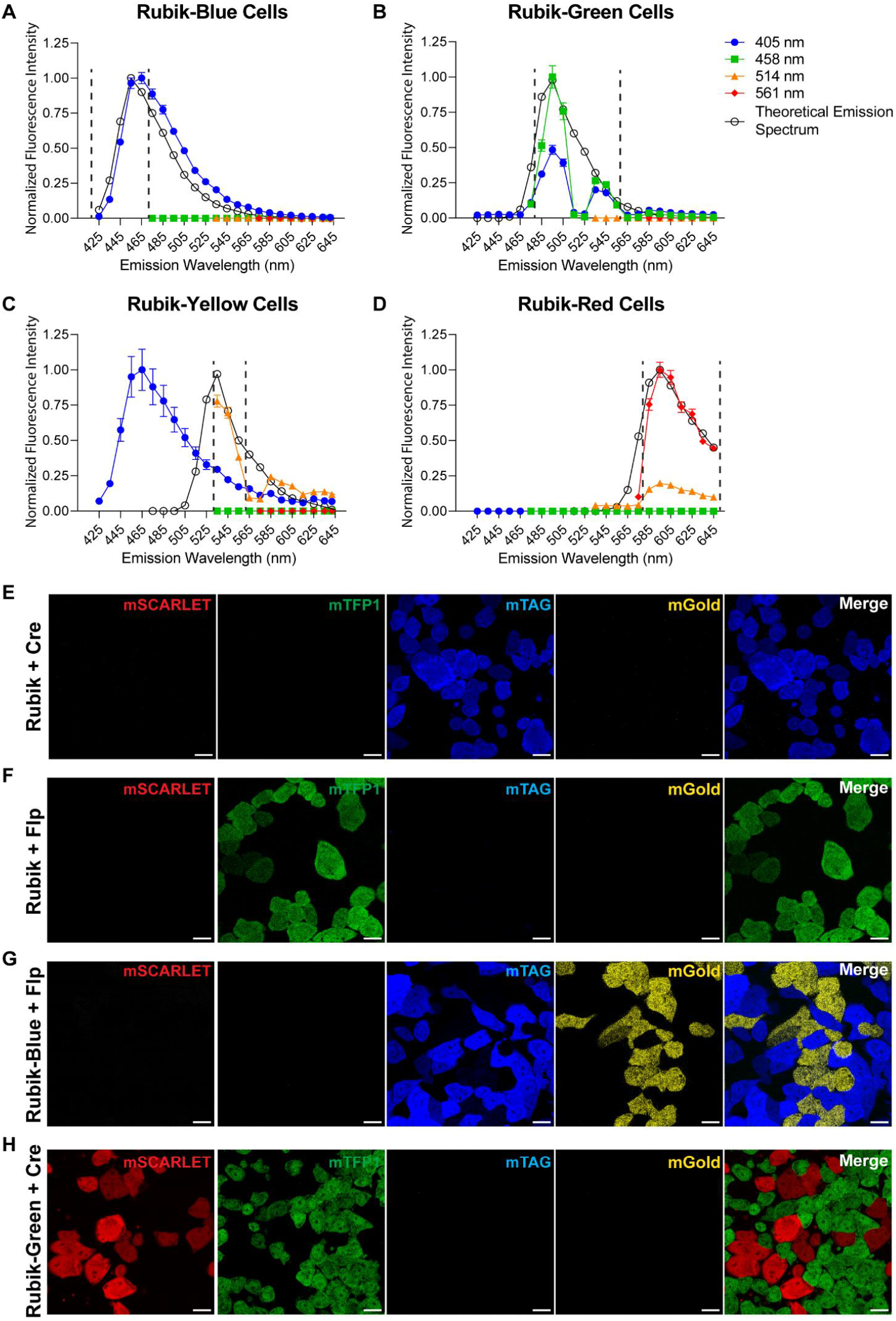
Fluorescence emission spectra and images to assess Rubik’s functioning in cell culture. (A-D) Fluorescence emission spectra of (A) Rubik-Blue, (B) Rubik-Green, (C) Rubik-Yellow, and (D) Rubik-Red cells were obtained using excitation lasers at 405, 458, 514, and 561 nm. The dotted vertical lines represent the emission intervals selected to acquire images of Rubik-cells. In each graph, the theoretical emission spectrum of the corresponding fluorophore is reported from FPbase (https://www.fpbase.org/spectra/). At least 20 cells for each color were analyzed. Values are reported as means ± SEM. (E-H) Representative confocal images of Rubik-cells after (E) single Cre recombination, (F) single Flp recombination, (G) sequential Cre→Flp recombination, and (H) sequential Flp→Cre recombination. Scale bar: 20 µm.

**Figure 3.**
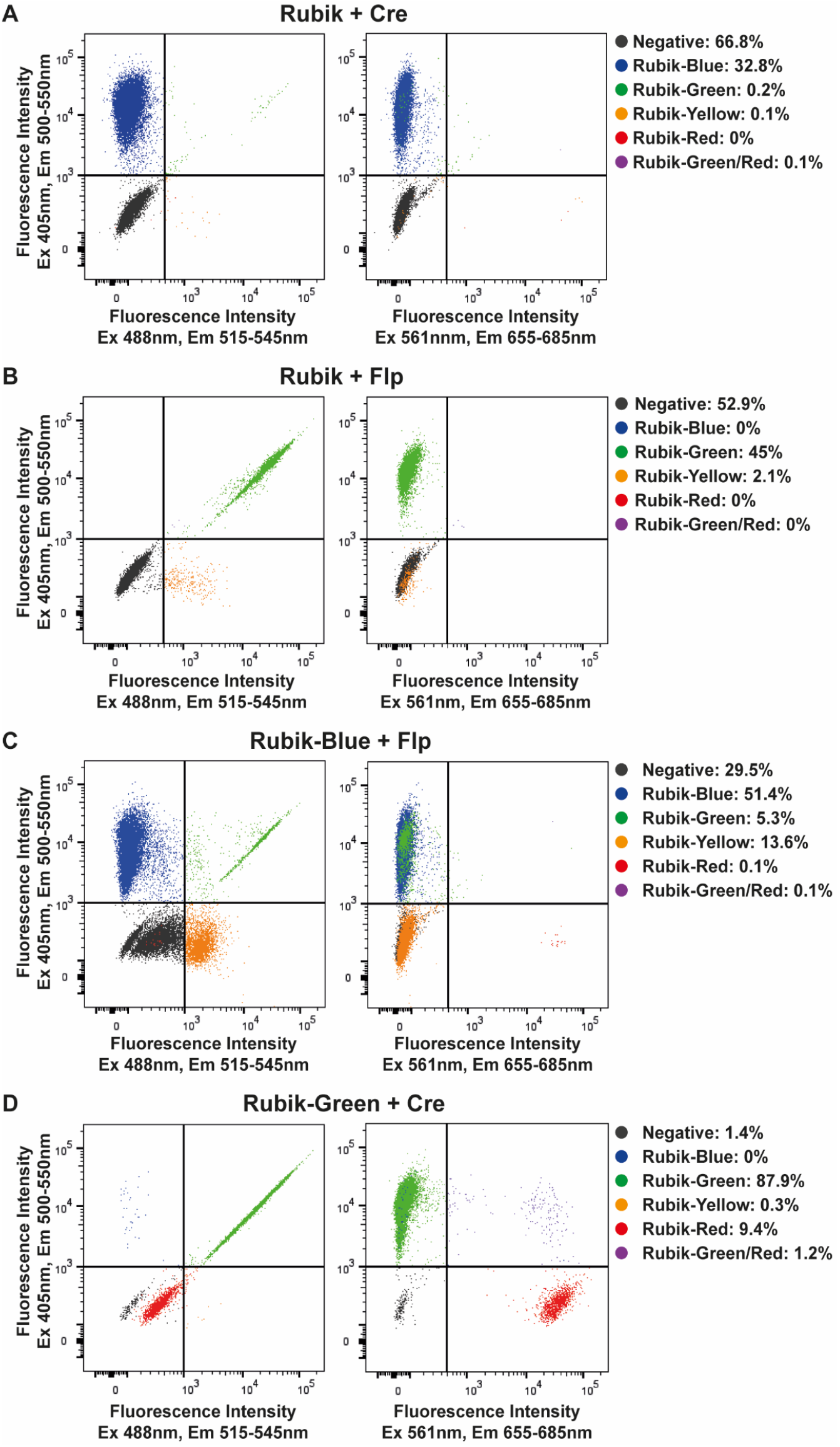
Flow cytometry plots to quantify Rubik’s recombination efficiency. Flow cytometry plots of Rubik-cells upon (A) single Cre recombination, (B) single Flp recombination, (C) Flp recombination of Rubik-Blue cells, and (D) Cre recombination of Rubik-Green cells. Graphs represent the fluorescence intensity detected with different combinations of excitation (Ex) and emission (Em) filters (Ex 405 nm and Em 500-550 nm; Ex 488 nm and Em 515-545 nm; Ex 561 nm and Em 655-685 nm). Quantifications of each cell population are reported as percentages.

### Rubik’s functioning with drug-inducible recombinases

To exploit Rubik’s full potential, it is crucial to exert tight temporal control over recombinase action. This can be achieved by having the recombination under the control of a double-inducible system. Therefore, we generated a trimethoprim-inducible optimized Flp recombinase (FlpO-DD) to use in combination with the tamoxifen-inducible Cre recombinase (ERT2CreERT2). We transfected Rubik-cells with either ERT2CreERT2 or FlpO-DD recombinases and administered tamoxifen (TAM) or trimethoprim (TMP) after 48 h, respectively. We then estimated the recombination efficiency by confocal microscopy and quantified it by flow cytometry. Both TAM and TMP treatments were able to effectively stimulate ERT2CreERT2 and FlpO-DD recombination, respectively (**Figure 4**). Among ERT2CreERT2-transfected cells, about 20% were mTagBFP2^+^ cells upon TAM treatment, while about 2% were mTagBFP2^+^ cells without TAM treatment (**Figure 4C**). Among FlpO-DD-transfected cells, about 33% were mTFP1^+^ cells upon TMP treatment, while about 10% were mTFP1^+^ without TMP treatment (**Figure 4D**). These results show that both inducible recombinases can be effectively activated by their respective drug, although for FlpO-DD a non-negligible degree of leakage (recombination activity without the drug) is observed.

**Figure 4.**
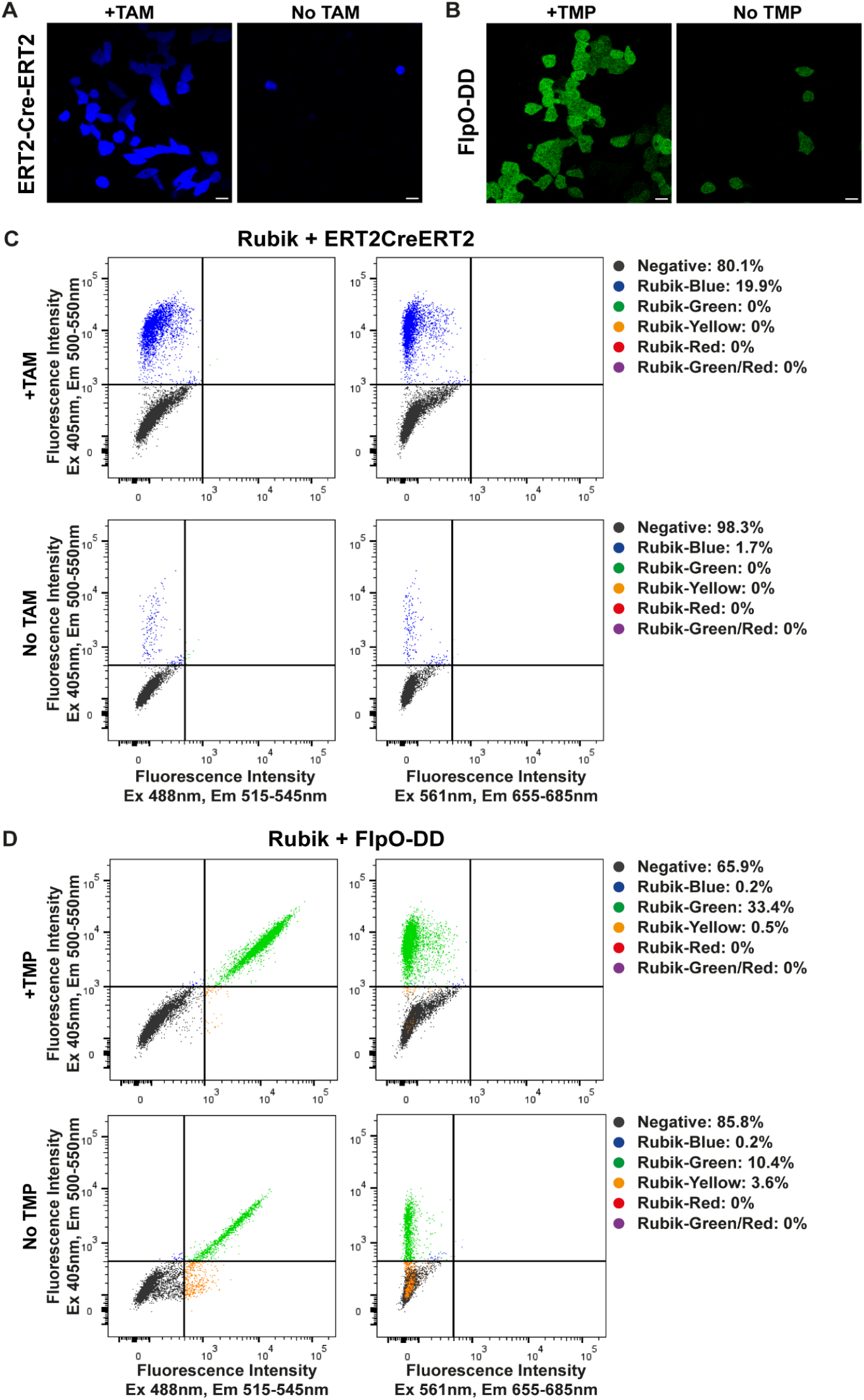
Rubik’s functioning with drug-inducible recombinases. (A-B) Representative confocal images of Rubik cells (A) transfected with ERT2CreERT2 and treated with/without TAM, (B) transfected with FlpO-DD and treated with/without TMP. Scale bar: 20 µm. (C-D) Flow cytometry plots of Rubik-cells (C) transfected with ERT2CreERT2 and treated with/without TAM, (B) transfected with FlpO-DD and treated with/without TMP. Graphs represent the fluorescence intensity detected with different combinations of excitation (Ex) and emission (Em) filters (Ex 405 nm, Em 500-550 nm; Ex 488 nm, Em 515-545 nm; Ex 561 nm, Em 655-685 nm). Quantifications of each cell population are reported as percentages.

## Discussion

Here we presented Rubik, a fluorescent reporter for Cre and Flp recombinase activity. Compared to the existing tools^11^, Rubik reports the single recombination events (Cre AND NOT Flp together with Flp AND NOT Cre Boolean operations), as well as their combination (Cre AND Flp Boolean operation) with the sequential information: Cre→Flp has a different outcome from Flp→Cre recombination. These two conditions are also associated with the co-expression of either excitatory or inhibitory optogenetic tools. Moreover, these light-gated ion channels have a tag to confine their expression to the soma for more precise spatial control of optogenetic stimulation. The possibility of having a single molecular tool to photo-excite or photo-inhibit identified neuronal populations is a unique feature that makes Rubik a highly versatile tool for neuroscience. Here, we showed the Rubik expected behavior in HeLa cells upon Cre and Flp activity and tested the correct DNA sequences after recombination. However, HeLa cells are not electrically excitable, and therefore we could not test the response to optical stimulation. This experiment would require neuronal cells expressing Rubik. Unfortunately, the Rubik size (about 10 kb) is not compatible with viral vector transduction, making tests in neurons difficult. Nevertheless, since the optogenetic proteins are under the control of the same promoter as the fluorescent proteins, and P2A and T2A are largely used and tested^16^, we can speculate that the opsins are expressed and functional. Moreover, since Rubik includes many recombination sites, the presence of multiple copies of the construct might result in unexpected recombination events. For this reason, we performed our experiments on single-clones of Rubik-cells generated using CRISPR-Cas9 technology. On the other hand, the best *in vivo* application of Rubik would be a transgenic mouse expressing the construct in a highly active transcriptional locus such as ROSA26^17^ or TIGRE^18^. Alternatively, Rubik could be transfected in specific brain areas using *in utero* electroporation. However, this approach would not address the multiple-copy problem, and proper recombination would need to be verified. To have better temporal control of Cre and Flp activity, we developed a double-inducible recombination system combining the existing TAM-inducible ERT2CreERT2 with an optimized TMP-inducible Flp recombinase (FlpO-DD). Indeed, the existing Flp-DD^15^ did not have sufficient activity in our experiments. The FlpO-DD showed a higher degree of leakage compared to the ERT2CreERT2 in HeLa cells. However, the leakage level and the drug dosage depend on the biological system and might vary across different cell types, such as neurons, and even more notably *in vivo*. Overall, our system offers, in a single molecular construct, a fluorescent reporter for Cre and Flp activity and an optogenetic tool to excite or inhibit identified neuronal networks. Moreover, by combining ERT2CreERT2/FlpO-DD with Rubik, photo-excitation or photo-inhibition can be achieved just by switching the order of drug administration. We think that Rubik represents a valuable resource for the neuroscience community since it would allow the identification and manipulation of neuronal subpopulations with an unprecedented degree of flexibility.

## Materials and Methods

### Rubik’s molecular structure

The Rubik plasmid was synthesized by GenScript. As a reporter, four different fluorescent proteins were chosen with minimal spectral overlap (see **Suppl. Figure 4**): mTagBFP2 (blue), mTFP1 (green), mGold (yellow), and mScarlet (red). A single CAG promoter was placed upstream of the four fluorescent protein genes, composed of the cytomegalovirus early enhancer element (C), the promoter elements of chicken β-actin gene (A), and the splice acceptor of the rabbit β-globin gene (G). The sequence of each fluorophore is followed by a polyadenylation signal, preventing the expression of more than one fluorescent protein at a time. The mGold fluorescent protein is fused with the soma-targeted channelrhodopsin-2 (ST-ChR2) through a P2A cleavage peptide, and the mScarlet fluorescent protein is fused with the soma-targeted Guillardia theta anion channelrhodopsin (ST-GtACR1) through a T2A cleavage peptide. Rubik’s sequence is flanked by two homology arms (HA-L and HA-R) to guide the Rubik insertion into the adeno-associated virus integration site 1 (AAVS1) locus on chromosome 19, a transcriptionally active site allowing a robust and stable expression of the transgene. Moreover, the system was equipped with three antibiotic resistance genes: (i) ampicillin resistance in the Rubik backbone for plasmid amplification; (ii) kanamycin/neomycin resistance after the HA-L site, allowing the selection of clones that have integrated Rubik in the right position; (iii) blasticidin resistance (BSD, blasticidin S deaminase) after the CAG promoter in the Rubik sequence to allow the selection of Rubik-expressing clones.

### Generation of a stable Rubik-cell line

HeLa cells were chosen because of their easy maintenance in culture and highly standardized genetic manipulation protocols. CRISPR-Cas9 technology was used to integrate Rubik into their genome. HeLa cells were cultured in DMEM (Gibco #31966021) supplemented with 1% penicillin-streptomycin (Gibco #15070063) and 10% Fetal Bovine Serum (FBS, Gibco #10270106) and transfected with Lipofectamine 3000 (ThermoFisher #L3000001) following the manufacturer’s protocol. To obtain a stable cell line with Rubik integrated, HeLa cells were double transfected with a plasmid containing the Cas9 nuclease and a sgRNA targeting the human AAVS1 locus (gift from Knut Woltjen; Addgene plasmid #80494; http://n2t.net/addgene:80494; RRID: Addgene_80494) and the Rubik plasmid. HeLa cells with Rubik integrated into the right locus were selected by adding geneticin (G418, Gibco #10131035) to the culture medium to a final concentration of 800 µg/ml, selected based on the antibiotic concentration that was able to kill 100% of WT HeLa cells over one week. After 10 days of antibiotic selection, HeLa cells with Rubik integrated into the transcriptionally active site AAVS1 (Rubik-cells) were obtained and plated using single-cell sorting (BD FACSymphony S6). After two weeks, single clones of Rubik-cells were obtained and used for the subsequent experiments.

### Cell transfection

Rubik-cells were transfected with Lipofectamine 3000 (ThermoFisher #L3000001) following the manufacturer’s protocol. Briefly, cells were plated on 24-well cell culture plates (BioFil #TCP011024) at a density to ensure about 80% confluency at the time of transfection. The DNA to lipofectamine ratio used was 1:3 (w/v). The P3000 reagent was used along with lipofectamine for all the transfections in a ratio of 1:2 (w/v) DNA:P3000. Typically, 0.5 μg of plasmid DNA was added to each well. The Cre and Flp plasmids were both obtained from restriction digestion of the pDIRE plasmid (gift from Rolf Zeller; Addgene plasmid #26745; http://n2t.net/addgene:26745; RRID: Addgene_26745). A volume of 1.5 μl of lipofectamine was diluted in 25 μl of serum-free medium Opti-MEM (Gibco #31985070). Lipofectamine was initiated by adding separately 0.5 μg of each plasmid vector diluted in 25 μl of Opti-MEM with added 1 μl of P3000 reagent. After 15 min of incubation at room temperature, the transfection mixture was added to the cell culture, and the cells were incubated at 37 °C and 5% CO_2_ for 24-48 h.

### Fluorescence microscopy images and spectral analyses

Cells were grown and transfected on polylysine (Sigma-Aldrich #P4832)-coated glass-bottom dishes (20 mm, Avantor #734-2906) and imaged using a 63x oil-immersion objective (Leica #11506429) on a Leica SP8 confocal microscope. During the imaging session, the culture medium was replaced with a Ringer’s solution (120 mM sodium chloride, 5 mM potassium chloride, 2 mM calcium chloride, 1 mM magnesium chloride, 25 mM sodium bicarbonate, 5.5 mM HEPES, 1.1 mM D-glucose, 10% fetal bovine serum, pH=7.4). To select the optimal combination of excitation and emission intervals for Rubik’s fluorophores with minimal spectral overlap, the emission spectra of mTagBFP2, mTFP1, mGold, and mScarlet in sorted Rubik-cells (see **Suppl. Figure 2** for the sorting strategy) were measured using the confocal microscope equipped with Acousto-Optical Beam Splitter (AOBS). For each condition, the fluorescence was excited with either 405, 458, 514, or 516 nm laser lines, and the emission was acquired in consecutive 10 nm wavelength intervals starting at 10 nm from the excitation and up to 650 nm. For each excitation wavelength, an emission spectrum was obtained by averaging the fluorescence intensity across at least 20 cells. The emission spectra were normalized on the max value among the four emission spectra, using Matlab software (Matlab R2023b, Natick, Massachusetts: The MathWorks Inc.). Plots were generated using GraphPad Prism software version 9 (GraphPad Software Inc., San Diego, CA, USA). The selected combinations of excitation and emission intervals to obtain Rubik-cell images were the following: for mTagBFP2, excitation at 405 nm and emission at 420-470 nm; for TFP1, excitation at 458 nm and emission at 480-560 nm; for mGold, excitation at 514 nm and emission at 530-560 nm; for mScarlet, excitation at 561 nm and emission at 580-650 nm. Rubik-cell images for each recombination setting were analyzed using these combinations of excitation and emission intervals, using ImageJ software (NIH, Bethesda, Maryland).

### Flow cytometry

Cells were detached from the culture plate using trypsin (Gibco #25200056) 0.05% for 5 min at 37 °C and pelleted by centrifugation for 5 min at 1100 x g. Cells were collected in polypropylene tubes (Sarstedt #55.526.006), and a volume of 500 μl of PBS 2 mM EDTA 0.5% BSA (buffer) was added to each tube. The tubes were centrifugated for 5 min at 1100 x g, the supernatant was discarded, and a washing step was performed by adding 500 μl of buffer to each tube. After vortexing and centrifugating the tubes as above, the cell pellet was resuspended using 200 μl of buffer. Data were acquired with BD FACSymphony S6 and analyzed with FlowJo software (Version 10, Ashland, OR: Becton, Dickinson and Company). Cell sorting experiments were performed in low-pressure conditions, using a 100 mm nozzle and a high-purity level sorting mask. Cells were sorted in two-step mode: first isolation of the selected population with a 4-way purity mask, followed by single-cell sorting with the ACDU (Automated Cell Deposing Unit) of the instrument, into a 96-well plate. Data were acquired with BD FACSymphony S6 and analyzed with FlowJo software.

### Genomic DNA extraction and PCR analyses

Sorted mTagBFP2^+^, mTFP1^+^, mGold^+^, and mScarlet^+^ cells were cultured for one week. Then, cells were dissociated with trypsin (Gibco #25200056) 0.05% for 5 min at 37 °C and pelleted by centrifugation for 5 min at 1100 x g. Cells were lysed in a buffer containing Tris HCl 100 µM, EDTA 1 µM, NaCl 1 M, SDS 0.2% (w/v) for 2 h at 57 °C, and DNA was isolated by isopropanol precipitation. PCR was performed using GoTaq® G2 DNA polymerase (Promega #M7841) following the manufacturer’s protocol. Primer sequences are reported in **Suppl. Table 2**. The Mastercycler Nexus X2 (Eppendorf) was used to run the PCR reaction.

### Cloning of the FlpO-DD plasmid

The sequence of the FlpO recombinase was amplified from the pDIRE plasmid (gift from Rolf Zeller; Addgene plasmid #26745; http://n2t.net/addgene:26745; RRID: Addgene_26745). The sequence of the destabilization domain (DD) was amplified from the 10XUAS-flp-DD plasmid (gift from Jing-W Wang; Addgene plasmid #124777; http://n2t.net/addgene:124777; RRID: Addgene_124777). The forward and reverse primers to amplify the FlpO recombinase were designed to include at the 5’ extremities the EcoRI and XmaI restriction sites, respectively. The forward and reverse primers to amplify the DD sequence were designed to include at the 5’ extremities the XmaI and NotI restriction sites, respectively (see **Suppl. Table 2**). PCR amplification was performed using NEBNext® High-Fidelity 2X PCR Master Mix (Neb #M0541) following the manufacturer’s protocol, using an annealing temperature of 62 °C. The Mastercycler Nexus X2 (Eppendorf) was used to run the PCR reaction. The PCR-amplified products were purified by agarose gel electrophoresis (0.8% agarose gel) and using the Wizard® SV Gel and PCR Clean-Up System (Promega #A9281). The purified FlpO fragment was digested with EcoRI and XmaI (Neb, EcoRI #R0101, XmaI #R0180), while the purified DD fragment was digested with XmaI and NotI (Neb, NotI-HF #R3189). The pCAG-ERT2CreERT2 backbone (gift from Connie Cepko; Addgene plasmid #13777; http://n2t.net/addgene:13777; RRID: Addgene_13777) was digested with EcoRI and NotI. The purified digested fragments were ligated using the T4 DNA ligase (Neb #M0202) in a ratio 1:3:3 (pCAG:FlpO:DD, w/w/w) following the manufacturer’s protocol. The ligation mix was transformed into JM109 Competent Cells (Promega #L2005) following the manufacturer’s protocol. The colonies were screened for the ligated plasmid by PCR using 5’FlpO_Forward and 3’DD_Reverse primers (see **Suppl. Table 2**). The Mastercycler Nexus X2 (Eppendorf) was used to run the PCR reaction. The plasmid DNA was purified using the PureYield™ Plasmid Midiprep System (Promega #A2492) following the manufacturer’s protocol.

### Treatment of Rubik-cells for inducible recombinases

Rubik-cells were transfected with either the TAM-inducible Cre recombinase (ERT2CreERT2) or the TMP-inducible FlpO recombinase (FlpO-DD). Tamoxifen (Sigma-Aldrich #T2859) or Trimethoprim (Sigma-Aldrich #T5648) were administered after 48 h at a final concentration of 1 μM. The efficiency of transfection and induction was assessed after 48 h.

