## Supplementary Material for "A sequential event-responsive fluorescent reporter for inducible Cre and Flp combinatorial recombination"

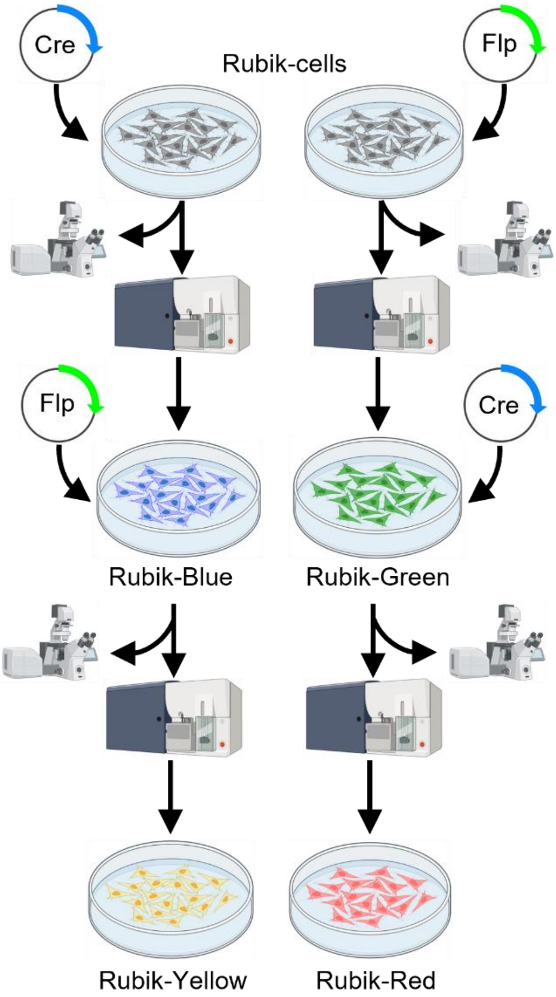


**Supplementary Figure 1. Schematic experimental protocol.** Schematic representation of the experimental steps to validate Rubik's functioning. Briefly, Rubik-cells were transfected with either Cre or Flp recombinases. The recombination efficiency was estimated by confocal microscopy and quantified by flow cytometry. mTagBFP2$^+$ (Rubik-Blue) and mTFP$^+$ (Rubik-Green) cells were isolated by FACS and transfected again with Flp or Cre recombinases, respectively. Finally, the recombination efficiency was estimated by confocal microscopy and quantified by flow cytometry.

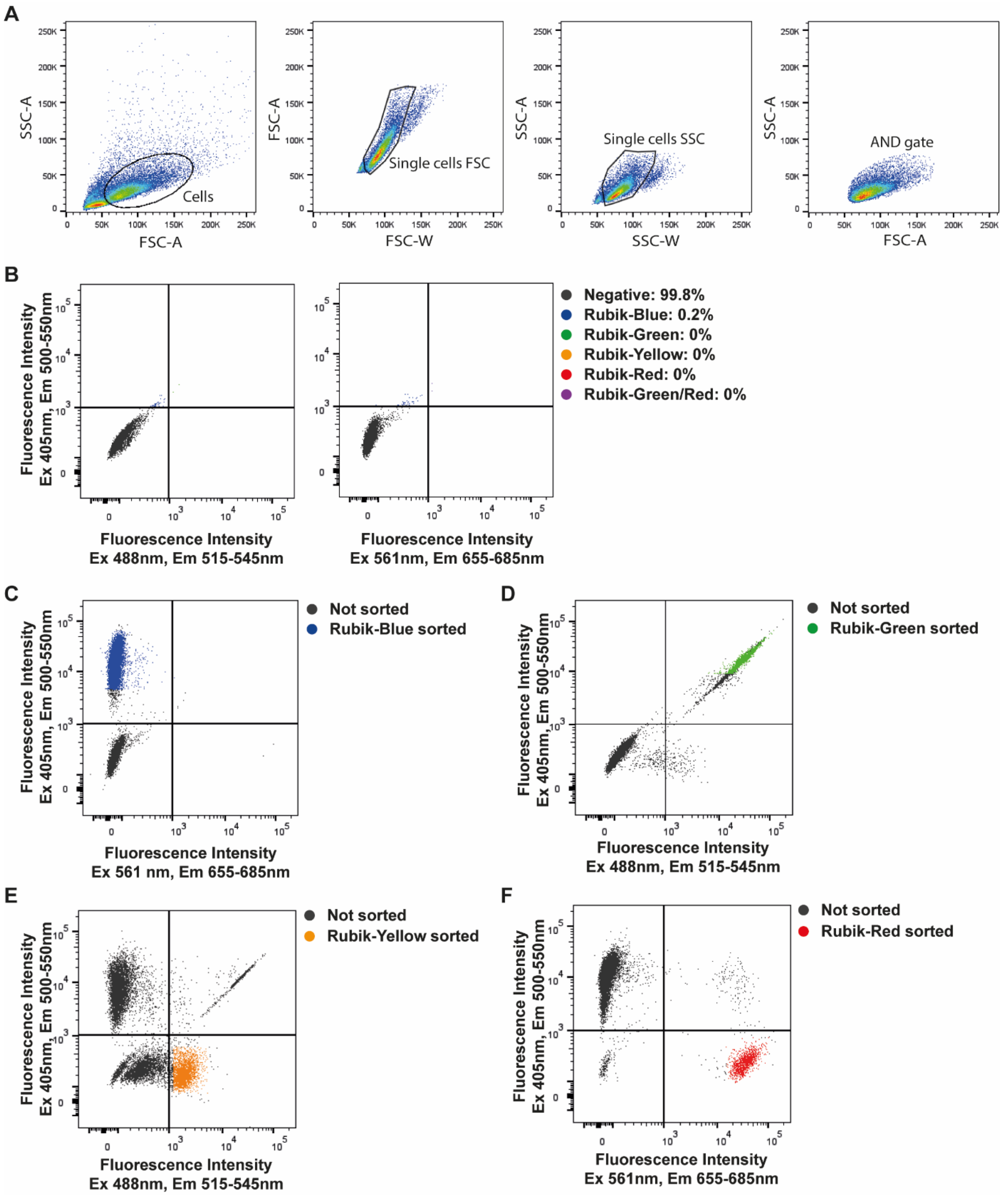


**Supplementary Figure 2. Gating and sorting strategy of Rubik-cells.** (A) Flow cytometry plots of Rubik-cells representing the strategy to determine the cell population by morphology and single-cell selection based on the side scatter (SSC) and forward scatter (FSC) area (A) and width (W). Cells are 52.1% of total events, single cells FSC are 91.8% of cells, and single cells SSC are 92.5% of cells. AND gate represents the Boolean logic operation between single cells FSC and SSC. Pseudocolors represent the increasing cell population density from blue to red. (B) Flow cytometry plots of non-transfected Rubik-cells used as a negative control. Graphs represent the fluorescent intensity detected with different combinations of excitation (Ex) and emission (Em) filters (Ex 405 nm and Em 500-550 nm; Ex 488 nm and Em 515-545 nm; Ex 561 nm and Em 655-685 nm). Quantifications of each cell population are reported as percentages. (C-F) Sorting strategy used to isolate (C) Rubik-Blue (93.2% of mTagBFP2$^{+}$ cells), (D) Rubik-Green (83.8% of mTFP$^{+}$ cells), (E) Rubik-Yellow (86.3% of mGold$^{+}$ cells), and (F) Rubik-Red cells (97.2% of mScarlet$^{+}$ cells). Colored dots represent sorted cells.

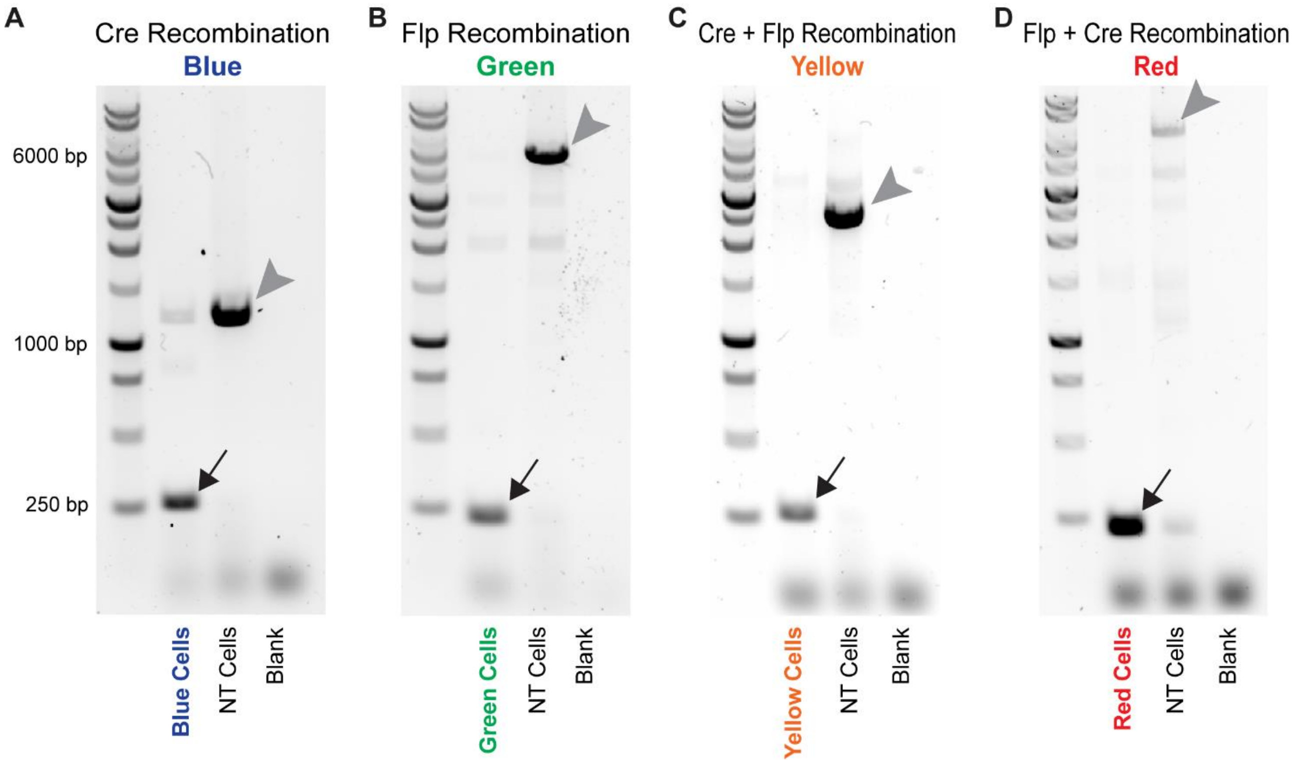


**Supplementary Figure 3. PCR validation of recombination events.** Agarose gel electrophoresis of PCR-amplified products using primer pairs specific for each recombination event. Black arrows indicate PCR bands corresponding to Rubik sequences after (A) single Cre recombination, (B) single Flp recombination, (C) sequential Cre→Flp recombination, and (D) sequential Flp→Cre recombination. Grey arrowheads indicate PCR bands corresponding to non-recombined Rubik sequences. NT = non-transfected cells.

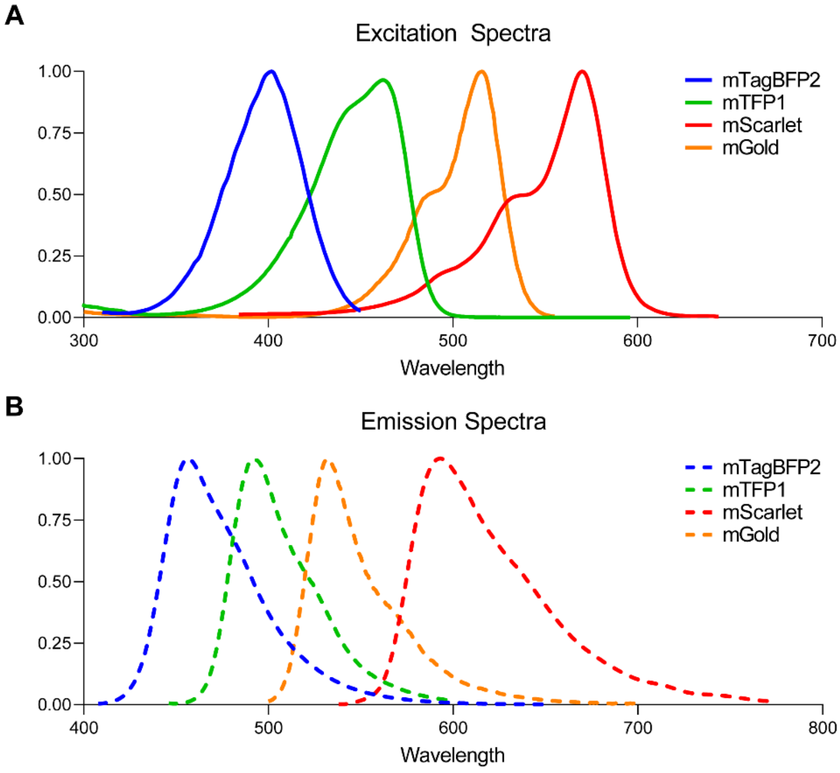


**Supplementary Figure 4. Excitation and emission spectra of the Rubik's fluorescent proteins.** Excitation (A) and emission (B) spectra of mTagBFP2, mTFP1, mGold, mScarlet fluorescent proteins from FPbase (https://www.fpbase.org/spectra/).

| Name | Sequence (5'→3') | Description |
|---|---|---|
| Rubik_Forward | GGGCAACGTGCTGGTTATTG | |
| Rubik-Blue_Reverse | ACGGTGCCCTCCATGTAAAG | Used together with Rubik_Forward to validate single Cre recombination. Expected amplicon sizes are 264 bp for the recombined sequence and 1227 bp for the non-recombined sequence. |
| Rubik-Green_Reverse | TACGCCCATTGTGGTCTCCT | Used together with Rubik_Forward to validate single Flp recombination. Expected amplicon sizes are 233 bp for the recombined sequence and 4808 bp for the non-recombined sequence. |
| Rubik-Yellow_Reverse | TAACACTGGTCCTCTGGCAC | Used together with Rubik_Forward to validate Cre→Flp recombination. Expected amplicon sizes are 263 bp for the recombined sequence and 2460 bp for the non-recombined sequence. |
| Rubik-Red_Reverse | ATAGATGGCTGGGTCGCAAG | Used together with Rubik_Forward to validate Flp→Cre recombination. Expected amplicon sizes are 230 bp for the recombined sequence and 6099 bp for the non-recombined sequence. |
| 5'FlpO_Forward | AAA**GAATTC**GCCACCATGGCTCCT | Used to amplify the FlpO sequence from the pDIRE plasmid. EcoRI and XmaI restriction sites are highlighted in bold. |
| 5'FlpO_Reverse | AAA**CCCGGG**GATCCGCCTGTTG | |
| 3'DD_Forward | AAA**CCCGGG**ATCTCCCTGATTGC | Used to amplify the DD sequence from the 10XUAS-flp-DD plasmid. XmaI and NotI restriction sites are highlighted in bold. |
| 3'DD_Reverse | AAA**GCGGCCGC**GATTCATTCTAGATT | |

**Supplementary Table 1. Primer sequences.** Primers used to validate Rubik recombination and to clone the FlpO-DD recombinase.
